# Accelerated Simulations of Chemical Reaction Systems using the Stochastic Simulation Algorithm on GPUs

**DOI:** 10.1101/2020.02.14.948612

**Authors:** James C. Pino, Martina Prugger, Alexander L. R. Lubbock, Leonard A. Harris, Carlos F. Lopez

## Abstract

Stochasticity due to fluctuations in chemical reactions can play important roles in cellular network-driven processes. Although the Stochastic Simulation Algorithm (SSA, aka Gillespie Algorithm) has long been accepted as a suitable method to solve the time-dependent chemical master equation, its computational cost is prohibitive for large scale complex networks such as those found in cellular processes. Here we present GPU-SSA, an implementation of the SSA formalism utilizing Graphics Processing Units for use in Python using the PySB modeling framework. We show that the GPU implementation of SSA can achieve significant speedup compared to parallel CPU or single-core CPU implementations. We further include supplementary didactic material to demonstrate how to incorporate GPU-SSA workflows for interested readers.

## 2 Introduction

Simulations of cellular processes at molecular resolution commonly rely on the chemical kinetics formalism to solve the Chemical Master Equation (CME). The most common approach is to represent chemical reactions using a system of Ordinary Differential Equations (ODEs), which is suitable for homogeneous mixtures. A key assumption of the ODE representation is that the underlying probability distribution of the chemical reactions must be unimodal [10]. For cells in particular, this is only obtained if the effective particle lifetimes (Kuramoto length) are sufficiently long and intermolecular distances sufficiently short to maintain the homogeneity and continuum assumptions [6]. Previous work has already highlighted the deviant effects that can emerge from ODE-based formalism [10]. Although these effects are often ignored for models that represent “average” cell-population mechanism, these effects may play important roles when considering the dynamics of cellular processes at the single-cell level [13].

Stochastic simulators of the CME, most notably the Stochastic Simulation Algorithm (SSA) formulated by Gillespie, provide a suitable solution to address the problem of stochasticity where the ODE-based formalisms are no longer applicable, such as single-cell processes [4, 5]]. From a computational perspective, the independence of each individual simulation in the SSA formalism makes parallelization an appealing approach for simulation speedup. Software packages such as StochKit [11] provide this form of parallelism using Central Processing Units (CPUs). In these implementations, the total simulation time decreases linearly with the number of CPU cores. However, for large and complex biochemical reaction systems, significant computational resources are required beyond typically available CPUs in a research setting. To address this limitation, researchers have increasingly turned to Graphical Processing Units (GPUs) to accelerate numerical simulations [2].

Previous implementations of the SSA algorithm in GPUs have already reported significant speedup, up to 200 times faster compared to CPU-based implementations [8]. However, these implementations have faced challenges associated with writing and maintaining specialized code in hardware-specific programming languages, as well as restricting the access of GPUs for novice users.

To address this challenge, we developed GPU_SSA, an implementation of the SSA solver in Python that can be used within the PySB [9] modeling framework developed in our lab, leveraging middle layer programming that allows minimal user involvement and easier code maintenance [7]. PySB enables users to encode biochemical reaction networks using a rule-based formalism in Python and supports import of models from other platforms including SBML [12]. We believe that our implementation will enable quantitative biologists, that lack specialized GPU knowledge, to carry out SSA-based simulations routinely. Our implementation includes two versions of GPU-SSA simulators: a CUDA version, specific to NVIDIA branded GPUs; and an OpenCL version that enables the use of parallel hardware independent of hardware platform. For a more thorough discussion about the implementation we refer the interested reader to the supplement.

## 3 Results and Discussion

To demonstrate the performance speedup of our GPU-SSA implementation, we compare our simulation run-times against the SSA solver included with BioNetGen [3] and the CPU-parallelized version of SSA provided by StochKit [11]. Details about the hardware and algorithms used to run the simulations can be found in the supplement.

Figure 1 shows the simulation output and timing results using the Extrinsic Apoptosis Reaction Model (EARM) [1]. Figure 1 top shows the output trajectories for cleaved cPARP accumulation upon apoptosis execution. We note that all SSA trajectories exhibit the snap-action switch behavior observed in apoptosis execution [1]. Figure 1 bottom shows the speedup from EARM simulations run with different implementations of the SSA formalism. For a large number of simulations, the single-processor BNG implementation of SSA runs for more than a week while the CPU parallel version of StochKit takes over 2 hours of wall-clock time using one processor. In contrast, the GPU implementation in GPU-SSA takes less than 20 minutes to evaluate all simulations. We note that the sequential run of StochKit was automatically optimized by the solver package at run-time, which explains the faster run-time compared to the BioNetGen solver. We also observe that CPU-based implementations of the SSA exhibit a linear dependence between number of simulations and run-time, i.e., doubling the amount of work also doubles the run-time. All models display a plateau at low number of simulations for the GPU-based SSA, suggesting that the problem size is too small to fully utilize the hardware until ~8,000 simulations. We observe a speedup of three orders of magnitude compared with the sequential BNG implementation for EARM. In the supplement, we present models with a speedup of four orders of magnitude. We attribute the differences in speedup between the OpenCL and the CUDA implementation of SSA to the fact that CUDA is specifically optimized to run on NVidia hardware.

**Figure 1:**
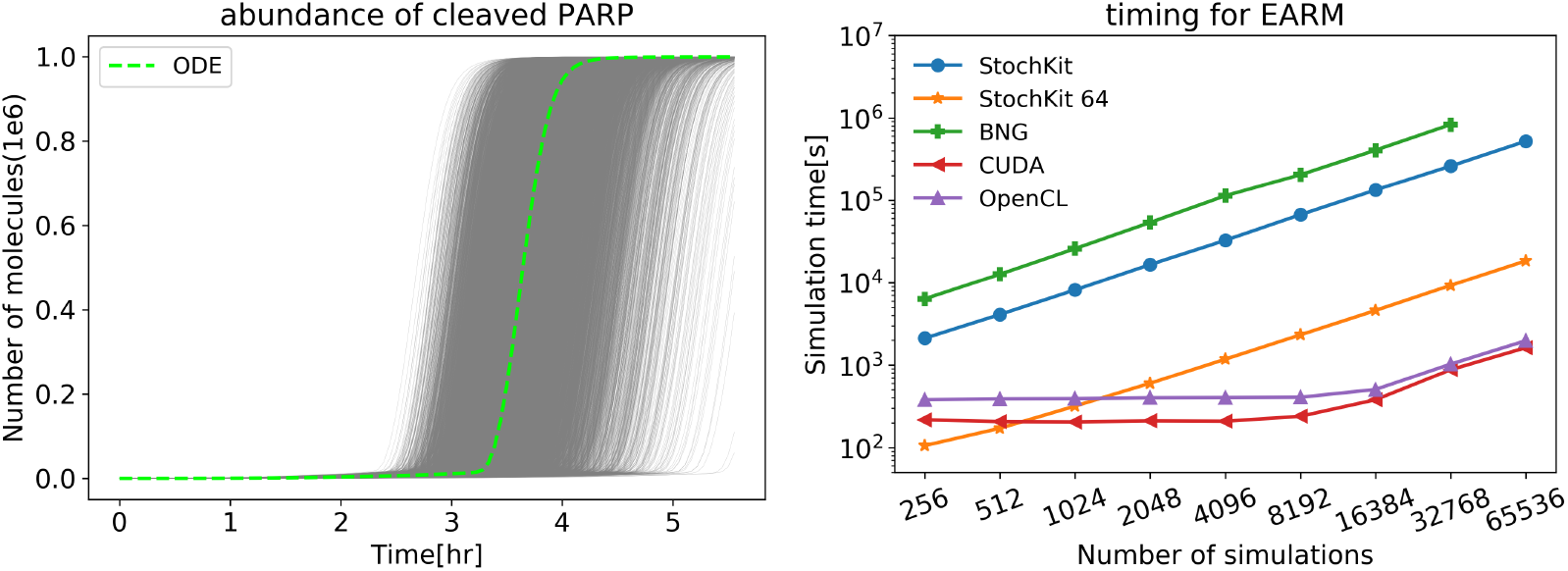
Top: Snap-action dynamics of PARP cleavage (cPARP) using the Extrinsic Apoptosis Reaction Model (EARM) [1] and GPU SSA. The gray lines represent individual SSA cPARP abundance trajectories while the dashed green line corresponds to the ODE-based solution. All simulations were run with the same simulation parameters. The observed response range thus emerged from intrinsic noise in the biochemical reactions. Bottom: Performance comparison for EARM SSA execution using BioNetGen, StochKit and GPU-SSA timings on the NVidia V100 hardware. BioNetGen (green) and Stochkit (blue) were run on one CPU. Stochkit was also run on 64 CPUs (orange). The GPU-SSA implementations in CUDA (red) and OpenCL (purple) were the fastest across all runs.

For small number of simulations (<1024), the parallel version of StochKit is faster than the GPU implementation, highlighting the overhead of GPUs for small processing loads. For large simulations, we observe that the resulting runtime on the GPU is an order of magnitude lower than the parallelized StochKit simulation on a single computational node. We provide timing results for various hardware including consumer-grade GPUs in the supplement.

In summary, we have implemented the SSA algorithm using available middle-ware packages in the Python programming language that will facilitate access to this simulator for all researchers. The GPU-SSA implementation is distributed using open-source licenses in PySB v. 2.0 and above. We invite the reader to read the supplementary documentation for further details.

