## supplementary information for "Accelerated Simulations of Chemical Reaction Systems using the Stochastic Simulation Algorithm on GPUs"

### Supplement

#### Contents

**1 Introduction** **1**

**2 Gillespies Direct Stochastic Simulation Algorithm** **2**

**3 GPU-SSA simulator: implementation and usage** **2**

**4 Validation of implementation** **3**

**5 Models** **6**

**7 Full tutorial using an SBML model from the BioModels database** **12**

#### 1 Introduction

Classical chemical kinetics (CCK) assumes a chemical system to be a continuum and can therefore describe the dynamics of the system using a system of deterministic Ordinary Differential Equations (ODEs). This approach, however, can only capture the mean behavior of the dynamics, and might underestimate the impact of low abundant species. Recent results from single-cell experiments have demonstrated that cellular processes can be highly heterogeneous, and thus challenge the traditional ODE-based approach to model cell-molecular processes. For example, cells treated with apoptosis-inducing drugs exhibit heterogeneous and non-deterministic response to treatment, underscoring the importance of stochasticity at the cellular level [7, 18]. Further examples for cell variability can be found, e.g., in [1, 5, 15]. Therefore, to study the molecular mechanisms that give rise to heterogeneous single-cell behavior, our simulation approaches must be reevaluated to better represent single-cell molecular processes.
Modeling the dynamics using stochastic chemical dynamics accounts for integer nature of molecular populations. Solving the Chemical Master Equation (CME) not only captures the behavior of low abundant species in the system, but also models the stochasticity that can take place in chemical reactions. Stochastic simulators of the CME, most notably the Stochastic Simulation Algorithm (SSA) formulated by Gillespie, provide a suitable solution to address the problem of stochasticity where the ODE-based formalisms are no longer applicable. Although this approximation does not explicitly deal with spatial heterogeneity, it does allow for the introduction of chemical reaction noise into the system, thus enabling researchers to explore the role of biochemical stochasticity in single-cell molecular processes.
Solving the CME using SSA requires a large number of simulations and therefore is highly dependent on computational power. Various optimizations have been proposed to reduce this cost by modifying the SSA algorithm, such as pre-sorting for reaction channels to speed up the determination of firing times of the direct method [3, 14], while other methods, such as tau-leaping [9] can trade speed for accuracy.
In this paper, we take an alternative approach to achieve feasible simulation times using state-of-the-art GPU hardware. Originally designed to quickly render computer graphics, modern GPUs are equipped with thousands of compute cores that can perform parallel calculations with high efficiency. Modern GPUs can provide the performance of a multi-CPU cluster in a small form factor that can be housed in a desktop workstation.

Utilizing the resources of modern GPUs, as well as the capacity of the programming language Python [19], we introduce a user-friendly implementation of the SSA direct method [8] to solve the CME, using the rule-based modeling framework PySB [13].

In section 2, we introduce the direct method of Gillespie that we used for our implementation. Section 3 explores the implementation of the algorithm on GPUs and how to use it within the PySB framework. In section 4, we validate our implementation by comparing it with the results of other software packages. Section 5 introduces the test models we used for our timing evaluation and the consequent timing results can be found in section 6. In section 7, we introduce a step-by-step tutorial using a model chosen from the BioModels database.

### 2 Gillespies Direct Stochastic Simulation Algorithm

For $\mathbf{N}(t) = (N_1(t), N_2(t), ..., N_n(t))$, where $N_i$ is the number of molecules of type $i$ and $n$ is the number of distinct chemical species, the chemical master equation (CME), given by

$$\frac{dP(\mathbf{N},t)}{dt} = \sum_{r \in \mathcal{R}} a_r\left(\mathbf{N} - \nu_r\right) P\left(\mathbf{N} - \nu_r, t\right) - \sum_{r \in \mathcal{R}} a_r(\mathbf{N}) P\left(\mathbf{N}, t\right),$$

describes the time evolution of the probability $P(\mathbf{N}, t)$ to be in a state $\mathbf{N}$ at time $t$. Let $\mathcal{R}$ be the set of all reactions occurring in the chemical system. In Gillespies direct method [8], the time development of each possible state depends on the reaction propensities $a_r(\mathbf{N})$, which are the probabilities per unit time, that reaction $r \in \mathcal{R}$ occurs given that the composition of the system is of the form $\mathbf{N}$. The rate of change is updated via the stoichiometric vector $\nu_r$, which describes the change in the numbers of each species as a result of the given reaction $r$.
The propensities $a_r(\mathbf{N})$ can be directly calculated via the reaction rate $\kappa_r$. For a first-order reaction of type $i$, the propensity is $a_r = \kappa_r N_i$. The second-order reaction of the same type $i$ has the propensity $a_r = \kappa_r N_i \left(N_i - 1\right)/2$, and for reactions of two different types $i$ and $j$ $a_r = \kappa_r N_i N_j$.
The Gillespie stochastic simulation algorithm (SSA) [8, 10] is simulating realizations of the state $P(\mathbf{N}, t)$ over time. Therefore, one simulation can be interpreted as one possible set of reaction events and the consequent changes in the molecular populations. To advance the simulation, we need to choose both a reaction time $\tau$ (which is exponentially distributed) as well as the reaction $\mu$ that fires at this time. The choice for reaction $\mu$ depends on the current state $\mathbf{N}$, as well as the reaction propensities (probabilities of a reaction happening per unit time), which depend on $\mathbf{N}$. After a reaction has fired, the propensities have to be updated, according to the new composition of $\mathbf{N}$. Algorithm 1 shows the basic step of the direct method.

initialization: set parameters, set $t = 0$
**while** *stopping criterion (exhaustion of a chemical, target simulation time reached, ...)* **do**
  For each reaction of type $i$, compute the propensities $a_i \forall\, i \in \{1, ..., M\}$ and $a_0 = \sum_{j=1}^{M} a_j$
  Generate two random numbers $r_1$ and $r_2$ in $\mathcal{U}(0,1)$
  Compute the random reaction time $\tau = \frac{1}{a_0} \ln\left(\frac{1}{r_1}\right)$
  Determine the next reaction by searching for the smallest integer $\mu$ that satisfies $\sum_{r=1}^{\mu} a_r > r_2 a_0$
  Update $\mathbf{N}$ according to the chosen reaction $r_\mu$
  Set $t \leftarrow t + \tau$
**end**

**Algorithm 1:** Stochastic simulation algorithm, direct method

### 3 GPU-SSA simulator: implementation and usage

Our parallelized version of the Gillespie Direct Method [8, 10] is part of the PySB Simulation Class [13]. Using PySB, the user can encode the desired chemical reactions of the biochemical system by representing reactions as interaction rules, or import already established models written in SBML code that are published on platforms such as the BioModels database. Our implementations leverage the PyCUDA and PyOpenCL Python packages [12], to generate and compile model-specific C code to be executed on the target hardware. Memory allocation (for parameters, initials values, and time courses) is automatically handled without the need for user input. Once compiled, the SSA algorithm is executed on the device and results are returned after all simulations have been completed. A schematic workflow is outlined in Figure 1.
To make use of this method, Python needs to be available on the computer and PySB needs to be installed. The PySB repositories as well as an installation introduction and tutorial for PySB can be found at `https://github.com/LoLab-VU/pysb`.

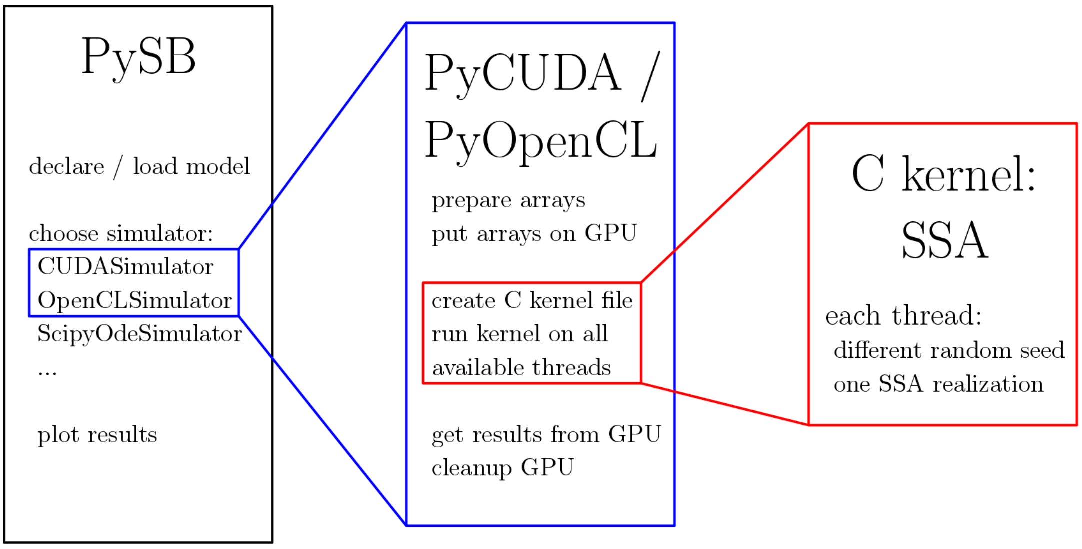

Figure 1: Workflow for the GPU-SSA implementation: a PySB model is imported and the GPU simulator is chosen. Both the PyCUDA version as well as the PyOpenCL version declare the working arrays for the SSA kernel, manage the memory allocation on the GPU, create the SSA kernel in C and pull the results from the SSA from the GPU back to the CPU. The kernel is created as a C-file that runs on every available hardware core. For each simulation, a thread is created that runs the full kernel, i.e. performs one realization of the stochastic simulation algorithm. After every thread is done with the simulation, the result vector is pulled back from the device and the device-memory gets cleaned up.

In Listing 1, we demonstrate a use case of the CUDASimulator for the Michaelis-Menten model provided with the PySB library.

Listing 1: Usage of the GPUSimulator in PySB.

```
#import the Michaelis−Menten model provided by PySB
from pysb.examples.michment import model
#import the CUDASimulator to solve the model
from pysb.simulators import CUDASimulator

# Set the simulation length and the number of time points
t = np.linspace(0, 20, 11)

# assign the CUDASimulator simulator to the model and run
sim = CUDASimulator(model)
# Run 1000 simulations
traj = sim.run(tspan=t, number_sim=1000)
```

For a step-by-step tutorial using an SBML model from the BioModels data base, see section 7.

#### 4 Validation of implementation

To validate our method, we use the Extrinsic Apoptosis Reaction Model (EARM) and compare it with the well-established SSA implementation provided by BioNetGen (BNG) [11]. Since SSA is a stochastic algorithm, comparing the results of GPU-SSA with BNG can only be done statistically. Using the standard parameters provided by the PySB implementation of EARM, we ran $10^3$ simulations up to a final time of $20^3$ seconds. This time frame is long enough for most simulations to cleave PARP, which is considered to be the point of no return in cell death. The observable proteins Bid, Smac, and PARP have been compared with each other.

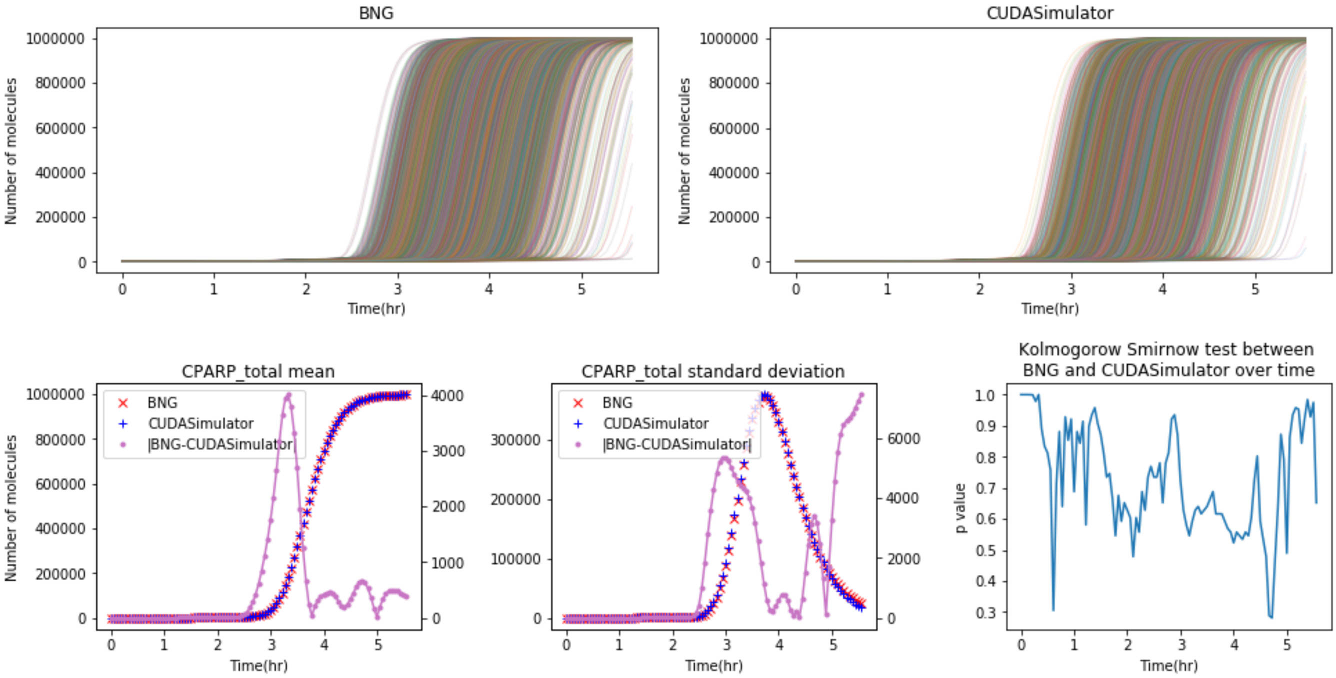

Figure 2: Time development of cleaved CPARP. On the top left hand side, we see the simulation performed with BioNetGen, on the top right hand side, we see the simulation performed with GPU-SSA. We observe that both methods need about 3-4 hours until the first simulation has all of its CPARP cleaved. The cleaving happens in a very rapid and short time frame. After 5.5 hours, most simulations have cleaved most of their CPARP and reached the point of no return for apoptosis. The main differences between the mean (Bottom left) and variance (Bottom middle) of the two methods occur during the time where CPARP starts to get cleaved. The scale on the left hand side denotes the abundance of CPARP and corresponds with the red line for the BNG simulation and the blue line for the CUDASimulator. The purple line denotes the absolute difference of each time point between the simulators and it scale can be found on the right hand side of the graph. We can see the largest difference in mean at the time where the cleaving of PARP is starting. However, the difference is about .4% of the total number of PARP in the system and can be explained by statistical variation. The Kolmogorov-Smirnov test (Bottom right) determines the smallest p-value of 0.279 at time point t=4.7 hours, which is far away from significant.

In Figure 2, we compare the mean and variance of the CUDASimulator with the BioNetGen simulator over time for the observable PARP. In red, we plot the mean and standard deviation of the BioNetGen simulation, in blue, we plot those modes for the GPU-SSA simulation. The purple line represents the total difference between the two methods. Both modes behave quite similar. The maximum difference in the mean of both methods occurs in the time frame between $10^3$ seconds and $15^3$ seconds, which is the time during which most of the simulations are starting their PARP cleavage. At the end time $20^3$, we observe, that nearly all PARP proteins are cleaved in both simulations. The behavior of the variance for both simulations is also very similar: we observe the maximum deviation within the simulation during the time frame at which most of the simulations have started the PARP cleavage.
The largest difference in the standard deviation can be found at the last time point. The Kolmogorov-Smirnov test is a measure whether two samples are drawn from the same distribution. A significant p-value would reject the hypothesis, that the samples come from two different distributions. The smallest p-value, however, is 0.279 at time point $t = 4.7$ hours, which is far away from significant. The data for GPU-SSA and BNG at each time step can therefore be considered drawn from the same underlying statistical distribution.

The observable Bid has a similar behavior to PARP, and we therefore omit the results at this place. The observable Smac, however, is another interesting case, since contrary to Bid and PARP, there is no smooth transition between the number of non-activated and activated proteins. After the permeabilization of the mitochondrial membrane (MOMP), Smac gets released at a single time point in a heavyside function. In Figure 3, we therefore include all the data for the release of Smac as well.

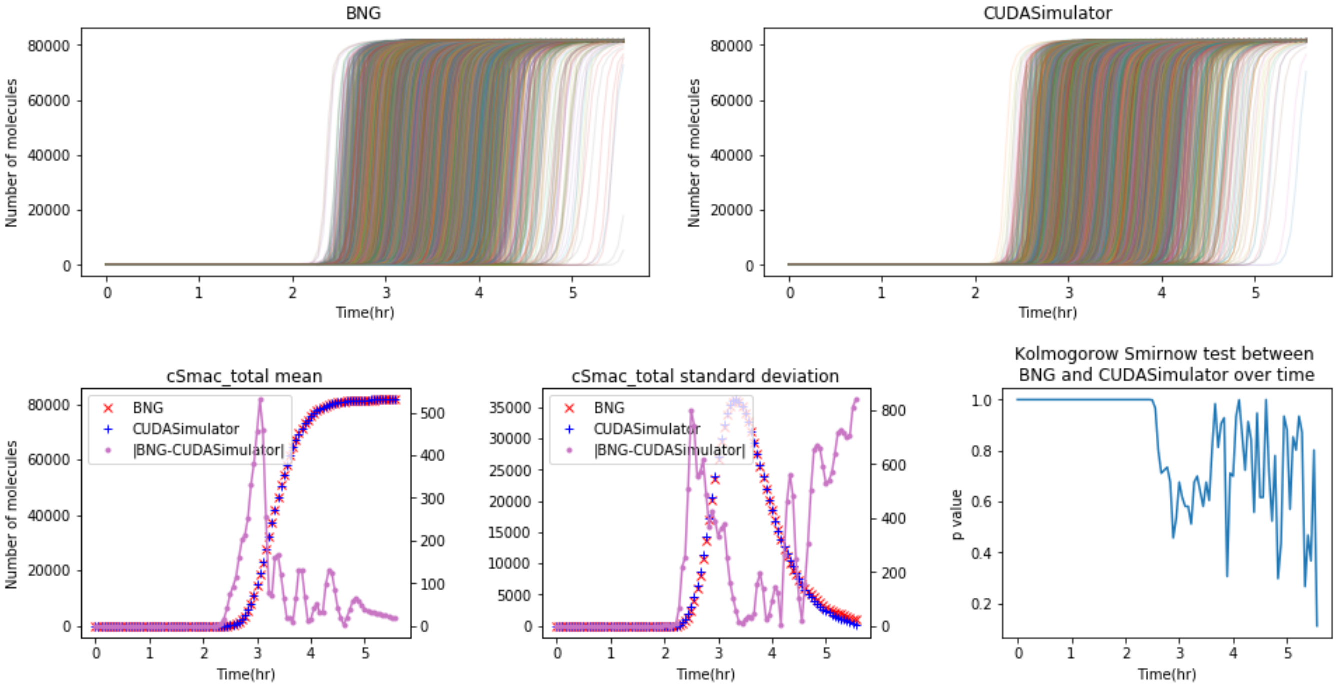

Figure 3: Time development of cleaved SMAC. On the top left hand side, we see the simulation performed with BioNetGen, on the top right hand side, we see the simulation performed with GPU-SSA. We observe that both methods need about 3-4 hours until the first simulation has all of its cSmac cleaved. The cleaving happens in a very rapid and short time frame. After 5.5 hours, most simulations have cleaved most of their SMAC and reached the point of no return for apoptosis. The main differences between the mean (Bottom left) and variance (Bottom middle) of the two methods occur during the time where Smac starts to get released. The scale on the left hand side denotes the abundance of cSmac and corresponds with the red line for the BNG simulation and the blue line for the CUDASimulator. The purple line denotes the absolute difference of each time point between the simulators and it scale can be found on the right hand side of the graph. We can see the largest difference in mean at the time where the cleaving of Smac is starting. However, the difference is about .6% of the total number of Smac in the system and can be explained by statistical variation. The Kolmogorov-Smirnov test (Bottom right) determines the smallest p-value of 0.11 at time point t=5.5 hours, which is far away from significant.

As we can see, the distributions are similar to each other. The smallest p-value in the Kolmogorov-Smirnov test is $p = 0.11$ at the last time point $t = 20000$. This is smaller than the smallest p-value for PARP, however it is still too large to be considered significant.

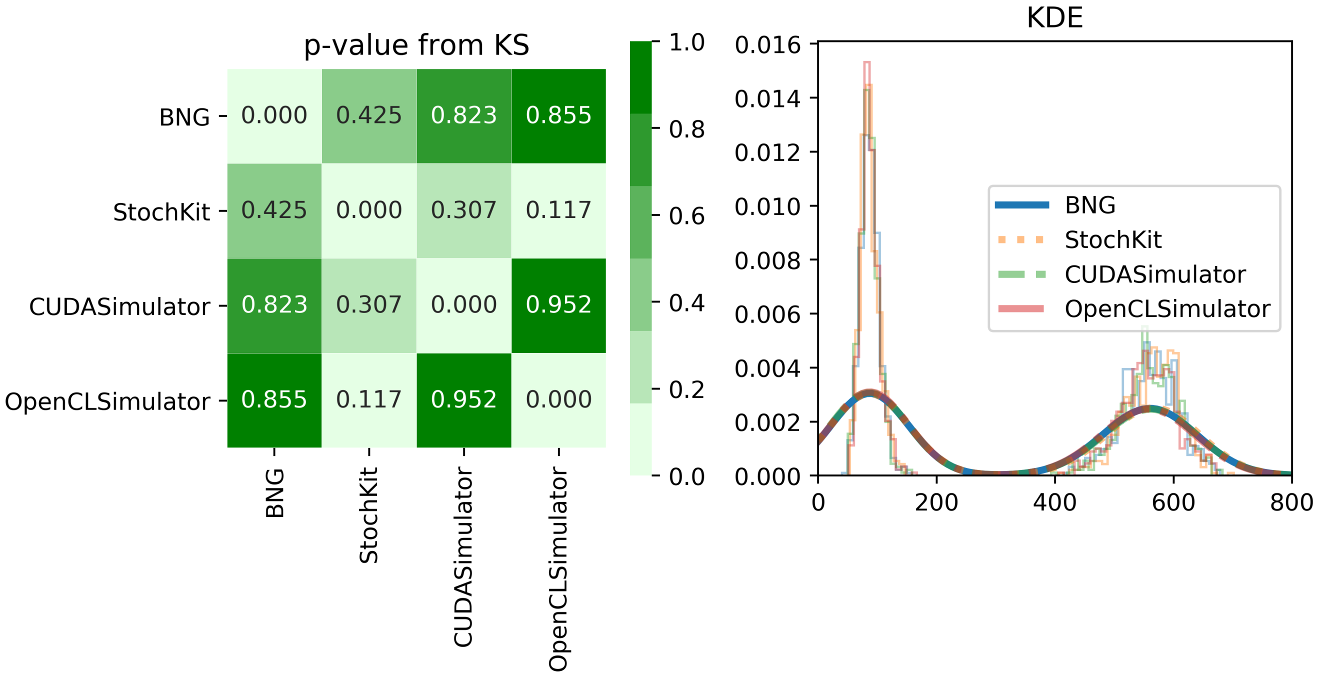

Figure 4: Validity check of the GPU implementations in comparison with the sequential BioNetGen SSA and the StochKit implementation. The check is performed using the Schlögl model and 1000 simulations. On the right hand side we see the end time of the simulation at which the check was done. The thin lines represent the histogram of each simulation, while the thick line represents the density plot of the same numbers. We can see that even though there are minor discrepancies in the histograms, they are averaged out in the density, such that we basically can not see any difference between the various methods at all. On the left hand side, we display the results of the Kolmogorov-Smirnov test taken at this time point. A significant p-value rejects the hypothesis, that the data was drawn from the same distribution. The smallest p-value in this case can be found between the comparison of the StochKit simulator and the OpenCLSimulator with a value of 0.117, which can not be considered significant. We therefore have no evidence to claim that any of those data is not drawn from the same distribution.

In Figure 4, we perform a full validity check over all simulators used in our study. For this check, we simulated the Schlögl model 1000 times with every simulator and compared the data points at a time, where all simulations reach the bi-stable steady state. The smallest p-value can be observed between the StochKit implementation and our OpenCLSimulator with 0.117, which can not be considered significant. We therefore conclude that there is no evidence that any of the simulation results in data point drawn from a different distribution.

### 5 Models

In this subsection, we introduce example models implemented in PySB and demonstrate the performance of our implementation versus BioNetGen and StockKit, the two SSA methods currently available to PySB users. All models are written in PySB and are available as examples on the github repository of PySB. The computational size of the models is summarized in Table 1.

#### 5.1 Michaelis-Menten

Michaelies-Menten kinetics can be seen as one of the most prominent test cases for biochemical reactions. It represents the reversible binding of an enzyme $E$ with a substrate $S$ to release a product $P$ in a non-reversible process. The process can be written as

$$E + S \underset{k_r}{\overset{k_f}{\rightleftharpoons}} ES \overset{k_c}{\rightarrow} E + P,$$

where $k_f$, $k_r$, $k_c$, denote the rate constants forward rate, reverse rate, and catalytic rate respectively.

| | Number of reactions | Number of species | Number of parameters |
|---|---|---|---|
| Michaelis-Menten | 3 | 4 | 4 |
| Schlögl | 4 | 3 | 6 |
| Kinase Cascade | 30 | 21 | 36 |
| EARM | 70 | 58 | 88 |

Table 1: Size of the different models used in this paper.

##### 5.2 Schlögl

In [17], Friedrich Schlögl introduced chemical reaction models that describe multi-modal steady states, a phenomenon at that time observed in thermodynamics. The chemical species in this example are representing rarefied gases and homogeneity is assumed due to constant stirring of the system.
One possible representation of such a non-equilibrium phase transition is

$$\begin{aligned} 2X &\underset{k_2}{\overset{k_1}{\rightleftharpoons}} 3X \\ I &\underset{k_4}{\overset{k_3}{\rightleftharpoons}} I + X. \end{aligned}$$

Note, that the concentration of $I$ remains constant over the simulation time, while the concentration of $X$ changes over time.
The Schlögl model is considered to be a standard test model to capture bi-stable states.

##### 5.3 Kinase Cascade

This model is adapted from [4].

##### 5.4 EARM

The extrinsic apoptosis reaction model (EARM) was introduced by [2] which describes the model pathway of apoptosis induced by an extrinsic signal. The system captures the permeabilization of the outer mitochondrial membrane (MOMP) resulting in the inhibition of apoptosis inhibitors, as well as the activation of apoptosome.

#### 6 Results

##### 6.1 Timing results

To demonstrate the performance speedup of our GPU-SSA implementation, we compare our simulation run-times against the SSA solver included with BioNetGen [6] and the CPU-parallelized version of SSA provided by StochKit [16]. For the StochKit simulations, we considered the performance of a sequential CPU run as well as parallelization across 64 cores. CUDA and OpenCL simulations were performed on a Volta V100 GPU while CPU simulations were performed on an IBM POWER9. To test the speedup relative to model size, we compare the four different biochemical models introduced in section 5. The models have a different degree of complexity, see Table 1. The Michaelis-Menten model and the Schlögl model [17] are similarly small problems. The latter model, however, is known to exhibit bi-stable states that are not captured by ODE-based solvers. The Kinase Cascade [4] is considered to be a medium-sized model, and EARM [2] is the largest model considered in our experiments. All models were encoded in PySB and are available in the examples folder of the PySB GitHub repository. The results of our experiments can be found in Figure 5.

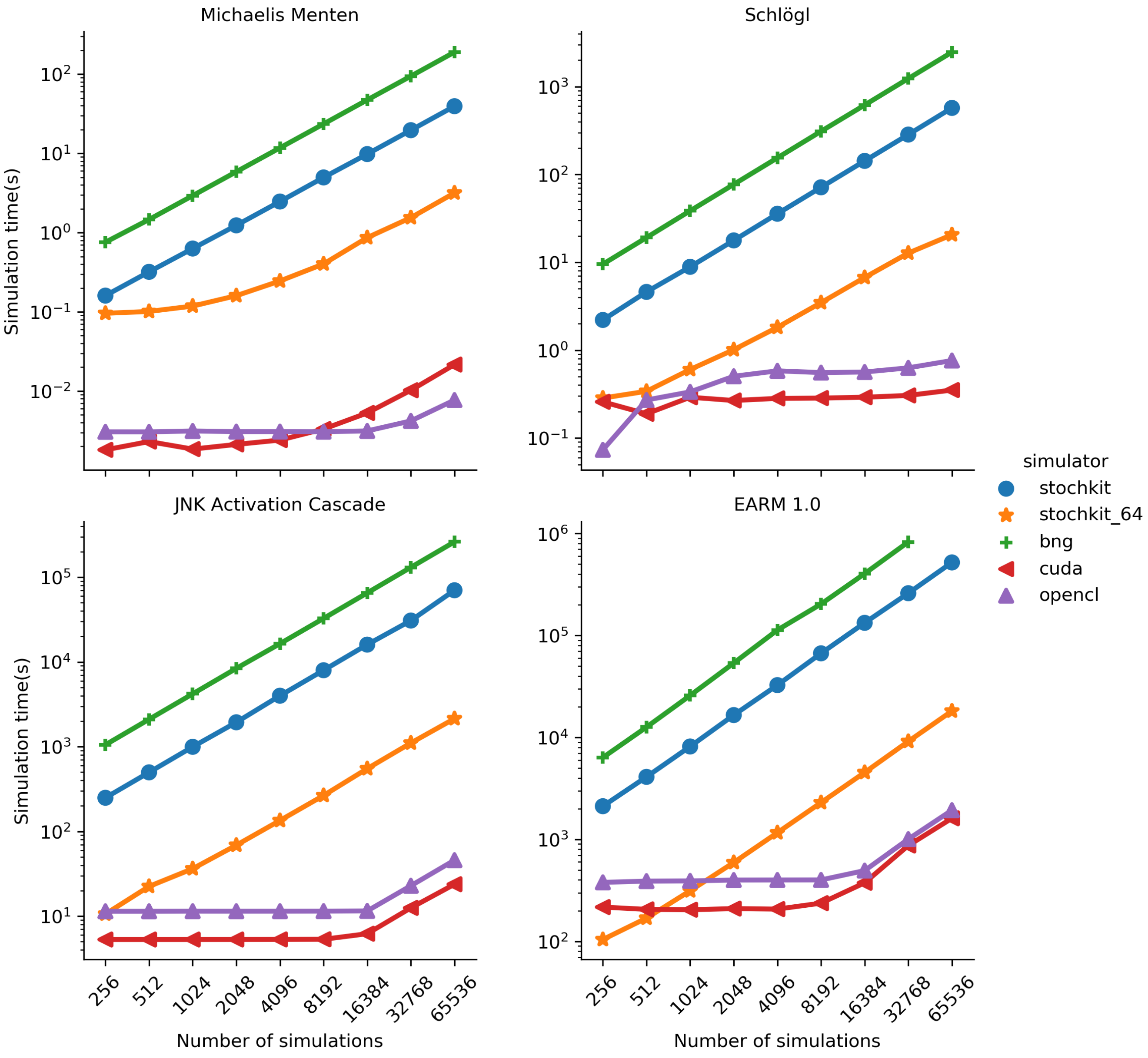

Figure 5: Comparing BioNetGen, StochKit and GPU-SSA timings run on the Nvidia V100. On the $x-$axis, we see the number of SSA simulations for each model. On the $y-$axis, we see the runtime for each simulation in seconds. Each simulation is a realization of a possible state of the system. BioNetGen depicted in green is implemented to run on only one CPU. In blue and orange, we represent the run-time for StochKit on 1 CPU, as well as a parallelization on 64 CPU cores, respectively.

On the $x-$axis, the number of simulations is increased by a factor of 2 for every model. On the $y-$axis, the runtime of the simulations is denoted in seconds. The BioNetGen (green) implementation of SSA can only run on one CPU core. As shown, doubling the number of simulations for each model also doubles the runtime for every model considered. This is also true for the sequential run of StochKit (blue). The smaller runtime for Stochkit compared to BNG hints at a more optimized baseline version of the algorithm. The orange line that describes the StochKit multi-CPU runtime has the potential to achieve increased speedup by using more CPUs. For example, in the EARM model we observe that for a small number of simulations, the parallel version of StochKit is faster than the GPU implementation on our system with 64 CPUs available. By adding more CPU cores to the problem, this line would be even lower. Note, however, that the Michaelis-Menten model only achieves a significant speedup from the CPU parallelization at a problem size of 4,000 simulations. This implies that the parallel overhead for such a problem size is more dominant than the actual simulation time. In this case, speedup by adding CPUs can only be expected for problems with more simulations. All models experience a plateau for the GPU simulators. This implies, that for the Michaelis-Menten problem ~8,000 simulations for the CUDA simulator and ~16,000 simulations for the OpenCL simulator are needed to fully exploit the hardware. The Kinase Cascade model fully utilizes the GPU for ∼16,000 simulations and EARM for ∼8,000 simulations. The continuing plateau in runtime of the Schlögl model suggests that the capacity of the GPU is not fully exploited until at least ∼65,000 simulations. For the small problems, the GPU solver experience about four orders of magnitude speedup compared to BNG and three orders of magntitude to the sequential StochKit run. The Kinase Cascade shows a speedup of 3.5 orders of magnitude for BNG and 3 orders of magnitude for StochKit. The EARM model experiences a speedup up to 3 and 2.5 orders of magnitude for the BNG and the sequential StochKit run respectively. The parallel version of StochKit on 64 cores beats the runtime of the GPU simulators for a small number of simulations for EARM, suggesting that the influence of the overhead of transferring data to and from the device is the main concern for a small work load. At ∼2,000 simulations, however, the simulation time is taking over the GPU overhead and starting at ∼16,000 simulations, the GPU implementations experience a speedup of an order of magnitude compared to the parallelized StochKit implementation. This speedup is even increased up to 2 orders of magnitude for thehttps://www.overleaf.com/project/5c6d9432950cbf5e22b90400 smaller models.

As we can see, both GPU implementations enable orders of magnitude faster simulations than the CPU implementations, even beating StochKit running across 64 CPUs. We attribute the differences in speedup between the OpenCL and the CUDA implementation of SSA to the fact that CUDA is specifically optimized to run on NVidia hardware.
In addition, we also include the fold change run time ratios in Figure 6 as well as the precise timing results in Table 2.

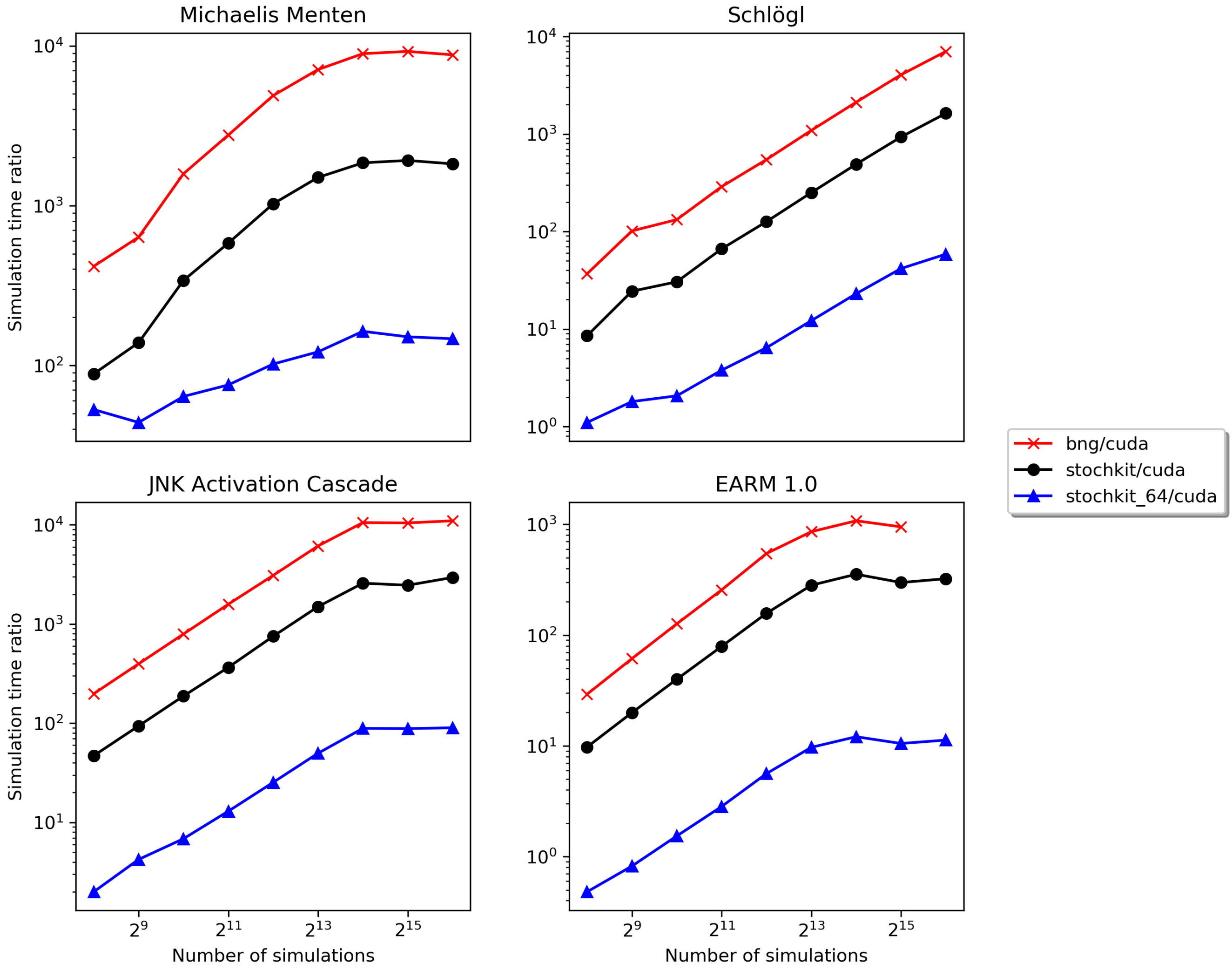

Figure 6: Ratio of run times between BNG, StochKit, StochKit on 64 cores, and CUDASimulator run on the Nvidia V100.

| model | n_sim | Simulator times (seconds) | | | | | Timing ratio | | |
|---|---|---|---|---|---|---|---|---|---|
| | | BNG | CUDA | OpenCL | StockKit | StochKit_64 | BNG \CUDA | StochKit \CUDA | StochKit64 \CUDA |
| **EARM 1.0** | **256** | 6348.13 | 216.74 | 378.81 | 2115.36 | 104.16 | 29.28 | 9.75 | 0.48 |
| | **512** | 12628.22 | 205.83 | 389.54 | 4104.66 | 168.92 | 61.35 | 19.94 | 0.82 |
| | **1024** | 25805.27 | 204.04 | 391.39 | 8150.94 | 313.07 | 126.47 | 39.94 | 1.53 |
| | **2048** | 53670.92 | 209.45 | 399.20 | 16577.10 | 593.39 | 256.24 | 79.14 | 2.83 |
| | **4096** | 113075.98 | 207.04 | 399.75 | 32606.80 | 1163.14 | 546.14 | 157.48 | 5.61 |
| | **8192** | 203555.37 | 236.80 | 400.28 | 66849.00 | 2296.05 | 859.58 | 282.29 | 9.69 |
| | **16384** | 404603.91 | 375.10 | 494.27 | 133109.00 | 4535.86 | 1078.65 | 354.86 | 12.09 |
| | **32768** | 827438.20 | 870.26 | 1005.73 | 259961.00 | 9144.92 | 950.78 | 298.71 | 10.50 |
| | **65536** | | 1610.57 | 1933.24 | 520748.00 | 18158.80 | | 323.32 | 11.27 |
| **Kinase Cascade** | **256** | 1051.37 | 5.29 | 11.41 | 249.35 | 10.58 | 198.47 | 47.07 | 1.99 |
| | **512** | 2102.81 | 5.29 | 11.41 | 497.49 | 22.44 | 396.98 | 93.92 | 4.23 |
| | **1024** | 4199.72 | 5.29 | 11.43 | 995.78 | 36.28 | 792.66 | 187.94 | 6.84 |
| | **2048** | 8393.11 | 5.30 | 11.43 | 1939.52 | 68.77 | 1583.29 | 365.87 | 12.97 |
| | **4096** | 16418.15 | 5.31 | 11.43 | 3993.20 | 134.53 | 3092.10 | 752.05 | 25.33 |
| | **8192** | 32634.33 | 5.33 | 11.45 | 7976.93 | 265.45 | 6122.02 | 1496.42 | 49.79 |
| | **16384** | 65257.74 | 6.20 | 11.51 | 16007.70 | 550.34 | 10524.15 | 2581.57 | 88.75 |
| | **32768** | 130748.75 | 12.51 | 22.84 | 30782.20 | 1104.34 | 10450.20 | 2460.29 | 88.26 |
| | **65536** | 261432.90 | 23.79 | 45.76 | 70123.80 | 2142.31 | 10986.69 | 2946.94 | 90.03 |
| **Michaelis Menten** | **256** | 0.755 | 0.002 | 0.003 | 0.160 | 0.096 | 417.16 | 88.42 | 52.88 |
| | **512** | 1.462 | 0.002 | 0.003 | 0.319 | 0.101 | 634.79 | 138.69 | 43.90 |
| | **1024** | 2.927 | 0.002 | 0.003 | 0.629 | 0.118 | 1581.74 | 340.03 | 63.85 |
| | **2048** | 5.858 | 0.002 | 0.003 | 1.234 | 0.160 | 2763.85 | 582.21 | 75.40 |
| | **4096** | 11.741 | 0.002 | 0.003 | 2.467 | 0.245 | 4881.25 | 1025.42 | 101.88 |
| | **8192** | 23.522 | 0.003 | 0.003 | 4.968 | 0.402 | 7102.32 | 1500.02 | 121.40 |
| | **16384** | 47.253 | 0.005 | 0.003 | 9.797 | 0.864 | 8932.86 | 1851.96 | 163.28 |
| | **32768** | 94.376 | 0.010 | 0.004 | 19.612 | 1.543 | 9226.00 | 1917.22 | 150.86 |
| | **65536** | 189.680 | 0.022 | 0.008 | 39.380 | 3.167 | 8786.72 | 1824.22 | 146.71 |
| **Schlögl** | **256** | 9.519 | 0.258 | 0.073 | 2.206 | 0.285 | 36.87 | 8.54 | 1.10 |
| | **512** | 19.118 | 0.188 | 0.271 | 4.602 | 0.340 | 101.47 | 24.42 | 1.80 |
| | **1024** | 38.555 | 0.291 | 0.335 | 8.881 | 0.602 | 132.52 | 30.52 | 2.06 |
| | **2048** | 77.159 | 0.267 | 0.504 | 17.755 | 1.012 | 288.67 | 66.42 | 3.78 |
| | **4096** | 154.493 | 0.282 | 0.581 | 35.702 | 1.819 | 547.08 | 126.42 | 6.44 |
| | **8192** | 310.031 | 0.285 | 0.555 | 71.408 | 3.472 | 1089.10 | 250.85 | 12.19 |
| | **16384** | 617.244 | 0.291 | 0.564 | 142.701 | 6.716 | 2119.73 | 490.06 | 23.06 |
| | **32768** | 1235.297 | 0.305 | 0.629 | 284.565 | 12.736 | 4049.66 | 932.88 | 41.75 |
| | **65536** | 2467.469 | 0.351 | 0.762 | 572.178 | 20.598 | 7025.57 | 1629.15 | 58.64 |

Table 2: Detailed timing results on the Nvidia V100 for the four models and the five different implementations. The numbers are in seconds. Due to the long runtime of the sequential BNG implementation, we did not run the 65536 simulations for this model.

#### 6.2 Timing comparisons of multiple GPU tiers

The simulations in this paper have been performed on a NVIDIA Volta GPU. For comparison, we also evaluated the K20C GPU as well as various generations of the less expensive GeForce consumer graphics cards (GeForce 980Ti, GTX1080, RTX2080). As shown in Figure 7, we saw improvements in timings across each generation of GPU. All GPU simulations were faster than the sequential CPU implementation for large number of simulations ($>$ 20k) on each machine (see Table 2). These data suggest that access to any recent GPU should prove to decrease simulation time compared to only using a CPU. We can also expect that future generations of GPUs will continuously increase the usefulness of our simulators.

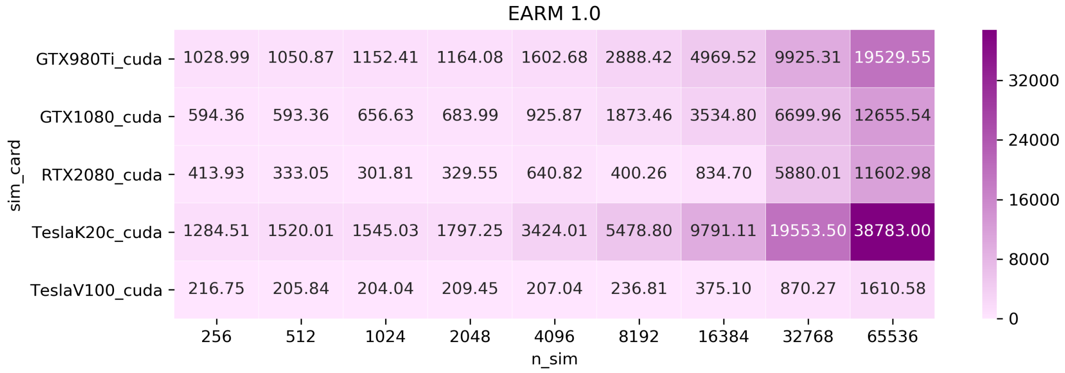

Figure 7: Timing analysis for various tiers of GPUs. From top to bottom, we benchmarked 5 GPUs (GeForce 980Ti, GTX1080, RTX2080, TeslaK20c, Tesla V100) across generations of graphics cards that we currently had access. We saw gradual improvement over the last three generations of consumer GPUs (980Ti, GTX1080, RTX2080). We included the timing of the Tesla K20c, a card released in 2012 (5 years prior to the Tesla V100), to demonstrate the dramatic increase in compute capabilities. We saw a substantial increase in performance of the Tesla V100 over the Tesla K20c, a 5 year difference in release dates. We also note that recent consumer grade cards continuously improve, yet are far behind compute centered cards.

#### 6.3 OpenCL

To make GPU-SSA more accessible to the user, we also offer an OpenCL implementation. Since CUDA is specific to Nvidia GPUs, this guarantees that the simulator is not vendor specific. We were unable to test the OpenCL implementation on the Volta GPU due to compatibility issues with our IBM CPU. Thus we used the NVIDIA RTX 2080 and a Ryzen 1700X CPU to demonstrate our OpenCL implementation. CUDA, OpenCL, and CPU timing for the four models is shown in Figure 8 .

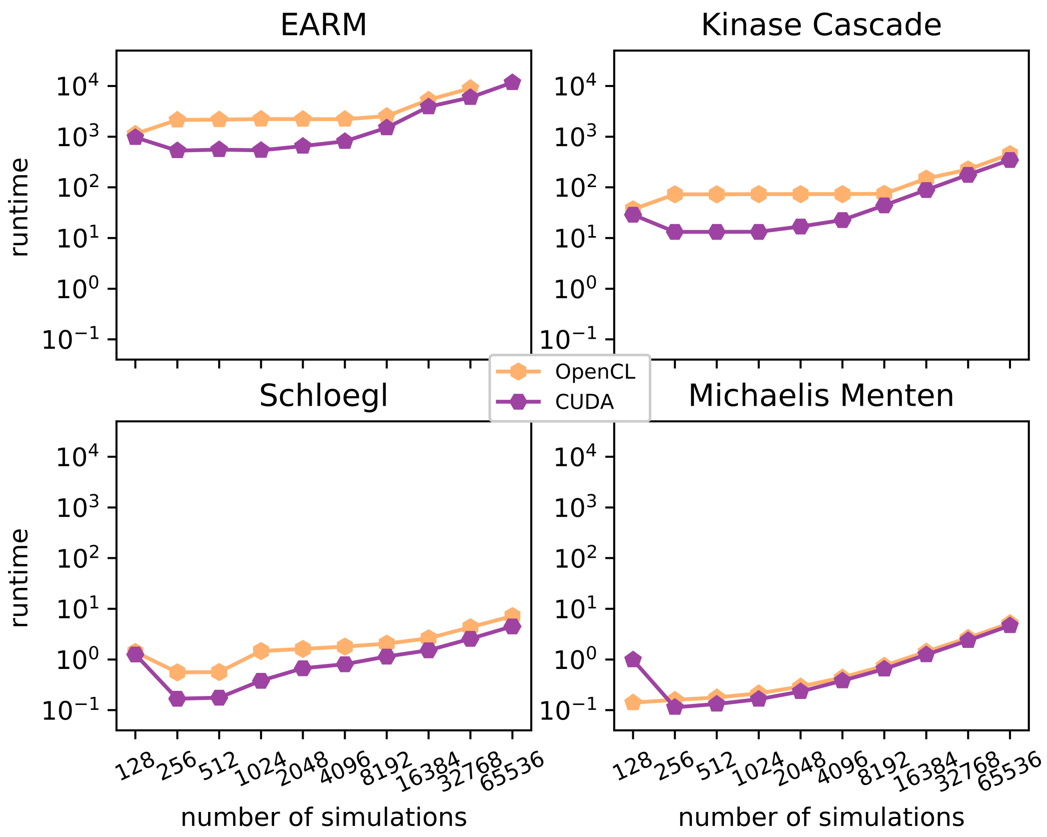

Figure 8: Timing comparison between CUDA and OpenCL. We can see the interplay of the Nvidia GPU with the CUDA simulator mostly for the simulations that have not yet reached the limits of the hardware. For all the models, we observe that when this limit is reached, the run time for both simulators increases similarly and the difference can only be observed for the relatively small Schlögl model. This leads us to our recommendation to use the CUDA simulator, if an Nvidia GPU is available. However, we are not too concerned about a possible OpenCL overhead.

Since CUDA is optimized for Nvidia GPUs, it is not surprising that the CUDA implementation runs more efficient than OpenCL. However, for EARM 1.0 and Kinase Cascade, we see the benefit getting less dominant when we have completely utilized the hardware. Schlögl shows some benefit from the CUDA implementation, while Michaelis-Menten, there hardly seems to be a difference between the two implementations at all. In principle, we would recommend using the CUDA implementation, if Nvidia GPUs are available. This analysis, however, makes us confident, that we do not loose too much performance using OpenCL instead, and that even non-Nvidia specific GPUs can give the user a significant benefit over a sequential implementation.

### 7 Full tutorial using an SBML model from the BioModels database

The GPU-SSA simulator is part of the PySB 2.0 (and above) simulation class. To access the solver with earlier releases, install the version directly from the GPU-SSA branch, as shown in Listing 2.

Listing 2: Requirements for the Jupyter Notebook tutorial.

```
#!/bin/bash

# Install miniconda from here: https://docs.conda.io/en/latest/miniconda.html

conda install numpy scipy cython sympy pandas
conda install -c alubbock bionetgen atomizer
pip install git+https://github.com/lolab-vu/pysb@gpu_ssa
pip install jupyter
```

Start Jupyter Notebook and create a new file in the Python interactive environment.

### Tutorial using an SBML model from the BioModels database

**Import libraries**

```
In [1]: import matplotlib.pyplot as plt
        import numpy as np
```

**Import the OpenCL version of the GPU-SSA implementation as well as the deterministic ODE solver**

```
In [2]: from pysb.simulator import ScipyOdeSimulator, OpenCLSimulator #CUDASimulator
```

**To be able to use an sbml-encoded model from the BioModels database, use the sbml importer from pysb**

```
In [3]: from pysb.importers import sbml
```

**Define the time points at which the model is evaluated**

```
In [4]: tspan=np.linspace(0,100,101)
```

**Due to the sbml importer from pysb, we can directly acces the available model from the BioModels database via the BioModels ID. In this case, the identifier BIOMD0000000008 stands for the cell cycle Goldbeter model by Gardner 1998 and models cell division cycle dynamics. We store the model from the database in the variable model.**

```
In [5]: model=sbml.model_from_biomodels('BIOMD0000000008')
```

**We run the SSA simulator for number_sim=100 simulations over the time span defined earlier. All resulting trajectories are stored in the variable traj. In this case we use the OpenCL version of the simulator. Since OpenCL can utilize various different hardware architectures, executing this step might result in the choice of available hardware on the host platform to be used for the simulation.**

```
In [6]: traj=OpenCLSimulator(model).run(tspan, number_sim=100)

Choose platform:
[0] <pyopencl.Platform 'NVIDIA CUDA' at 0x55616cfe7c40>
[1] <pyopencl.Platform 'Intel(R) OpenCL HD Graphics' at 0x55616ce8cfb0>
Choice [0]:0
Set the environment variable PYOPENCL_CTX='0' to avoid being asked again.
```

For comparison, the deterministic solution of the model is computed via the Scipy simulator for ODEs provided by PySB and stored in the variable y.

```
In [7]: y=ScipyOdeSimulator(model, tspan=tspan, compiler='cython').run()
```

Every model defines its own observables. Observables in this particular BioModel are cyclin_Cell, protease_Cell, cdc2k_Cell, cyclininhibitor_Cell, and complexinhibitor_cyclin_cell. The loop produces the same plot separately for each observable. The trajectories for the selected observable are unstacked into the x variable which aren then plotted with a reduced alpha value to reduce the dominance of 100 lines in the plot. The red and blue line outline the maximum and minimum value of the SSA simulation respectively. The black solid line depicts the mean of the SSA simulations. The green line, in contrast, is the result of the deterministic ODE

```
In [8]: for i in model.observables:
            x=traj.dataframe[i.name].unstack(0).values
            plt.figure()
            plt.title(i.name)
            plt.plot(tspan, x,'0.5', lw=2, alpha=0.25)# individual trajectories
            plt.plot(tspan, x.mean(1),'k-*', lw=3, label="Mean")
            plt.plot(tspan, x.min(1),'b--', lw=3, label="Minimum")
            plt.plot(tspan, x.max(1),'r--', lw=3, label="Maximum")
            plt.plot(tspan, y.dataframe[i.name],color='lime', \
                     linestyle='dashed', lw=3, label="ODE")
            plt.xlabel('Time')
```

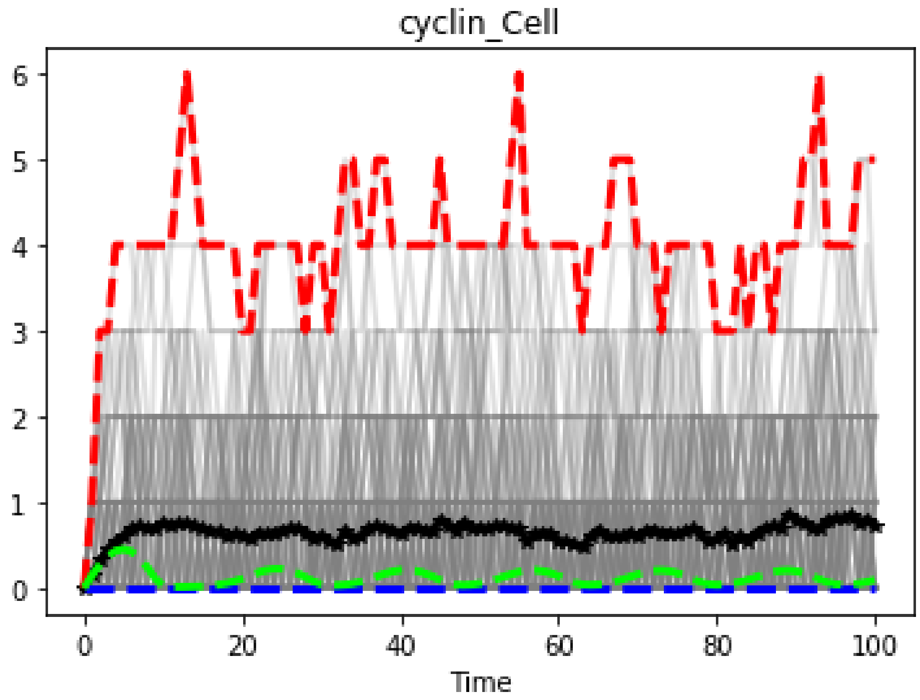

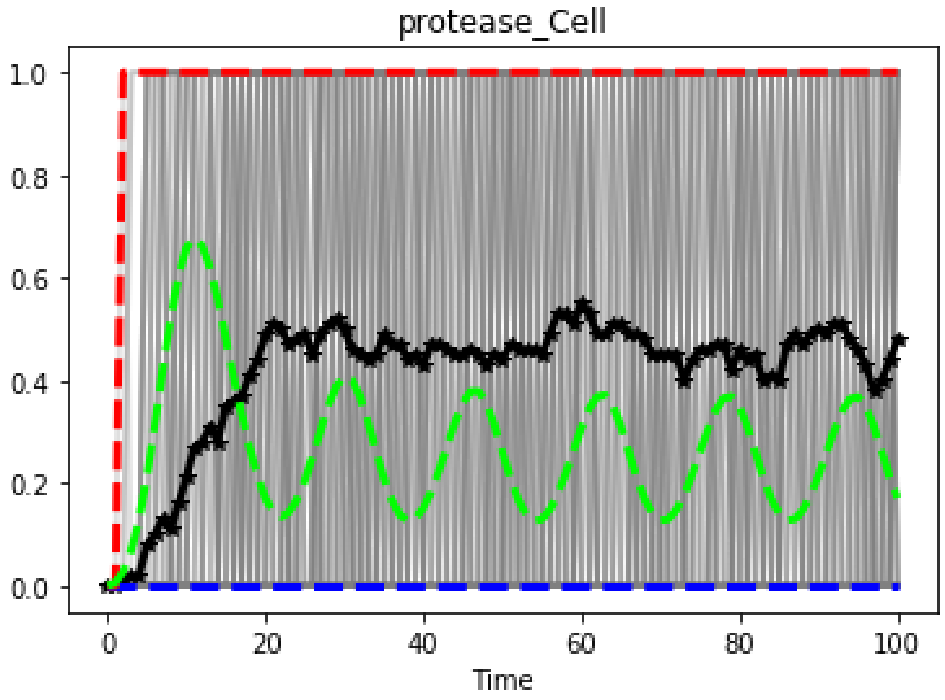
protease_Cell
1.0
0.8
0.6
0.4
0.2
0.0
0
20
40
60
80
100
Time

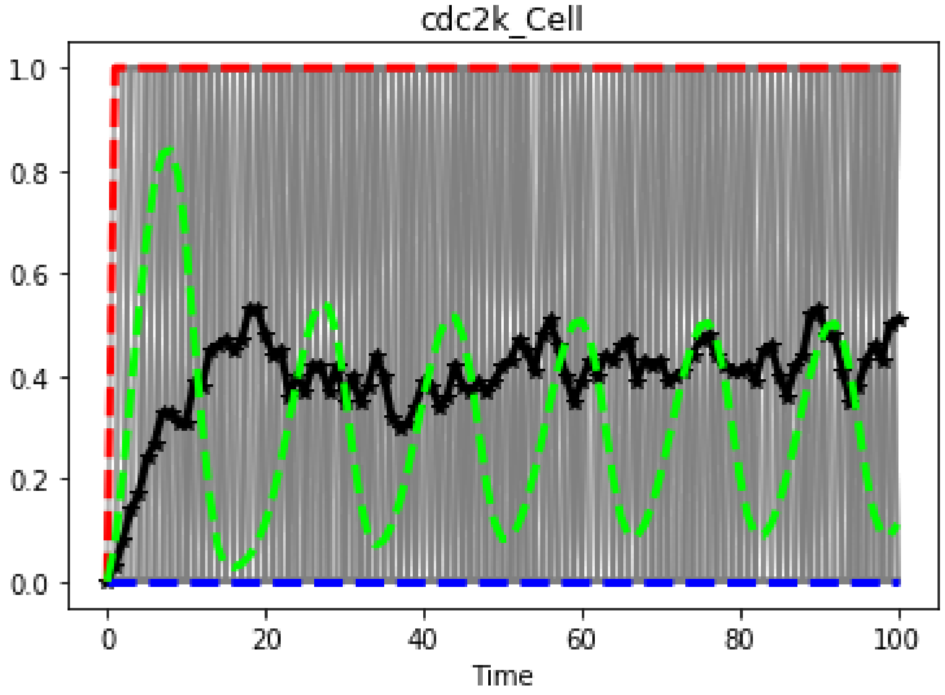
cdc2k_Cell
1.0
0.8
0.6
0.4
0.2
0.0
0
20
40
60
80
100
Time

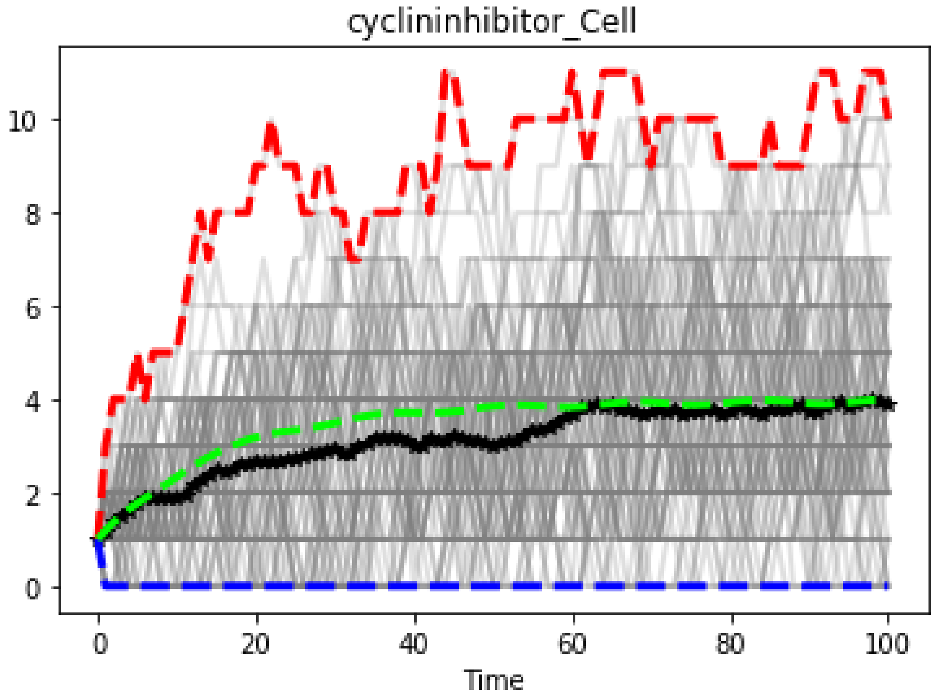
cyclininhibitor_Cell
10
8
6
4
2
0
0
20
40
60
80
100
Time

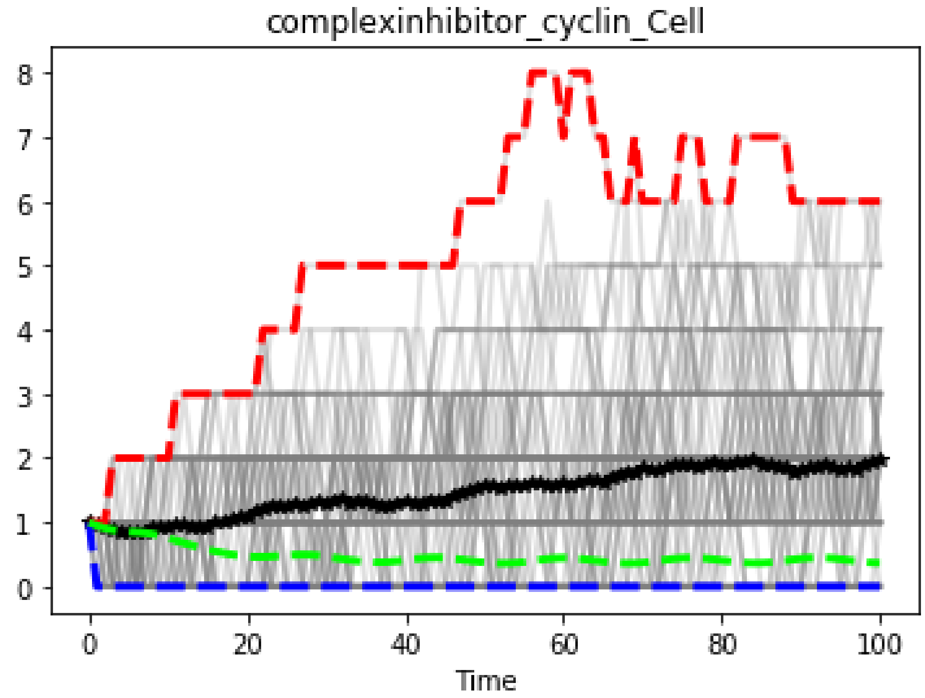
complexinhibitor_cyclin_Cell
8
7
6
5
4
3
2
1
0
0
20
40
60
80
100
Time
